# Comparison of CRISPR-Cas-based knockdown of endogenous mRNA in sensory neurons

**DOI:** 10.64898/2026.02.06.704510

**Authors:** Andrea Meulenberg, Macarena Pavez, Emma K. Gowing, David Mayo-Muñoz, Nils Birkholz, Graceallah Suhono, Peter C. Fineran, Robert. D. Fagerlund, Laura F. Gumy

**Author notes:** These authors contributed equally (Andrea Meulenberg, Macarena Pavez). DMM; Department of Plant and Environmental Science, University of Copenhagen, Frederiksberg, Denmark.

## Abstract

RNA-targeting CRISPR-Cas systems enable modulation of gene expression without permanent genome modification, making them useful for sensitive cell types such as neurons. While CRISPR-Cas technologies have been most extensively applied and validated in primary hippocampal and cortical neurons, their use in sensory neurons remains largely unexplored. Sensory neurons are an established cellular model for studying axon growth and regeneration, pain mechanisms, sensory transduction, and neuron–environment interactions. Here, we evaluated the performance of compact RNA-targeting CRISPR-Cas effectors Cas7-11S, hfCas13X, and hfCas13d in primary rat sensory neurons in culture. Using an endogenous mRNA as the target, we compared knockdown efficiency and assessed the effects of CRISPR-Cas expression on neuronal health. The systems showed distinct differences in performance, with Cas7-11S inducing toxicity, hfCas13X showing minimal knockdown, and hfCas13d providing robust gene silencing with minimal adverse effects on neuronal health. These findings identify hfCas13d as an effective and well-tolerated RNA-targeting CRISPR-Cas tool for sensory neurons and provide important insight into its suitability for neuroscience research and potential therapeutic applications.

## Introduction

Clustered regularly interspaced short palindromic repeats (CRISPR) and their CRISPR-associated (Cas) proteins are adaptive immune systems used by archaea and bacteria to defend against foreign DNA and viruses (Amitai & Sorek, 2016; Sternberg et al., 2016). These systems have been adapted into widely used tools for gene manipulation in basic research and therapeutic development (Abudayyeh et al., 2017; Becirovic, 2022; Wright et al., 2016). While DNA-targeting CRISPR-Cas tools are commonly used for genome editing (Cong et al., 2013), permanent modification of the genome is not always desirable. This is particularly true for therapeutic applications, where gene regulation may need to be reversible, and for cell types that are sensitive to DNA damage. RNA-targeting CRISPR-Cas systems provide a powerful alternative approach by enabling modulation of gene expression without altering the DNA sequence, making them more suitable in certain biological contexts (Zhu et al., 2024).

Neurons are a particularly challenging cell type in which to apply gene-modulating tools. As post-mitotic cells, they rely on stable and tightly controlled gene expression to maintain function, and their highly polarized morphology and dependence on long-distance intracellular transport make them especially sensitive to cellular stress (Gumy & Hoogenraad, 2018; Karra & Dahm, 2010; Zeitelhofer et al., 2007). While CRISPR-Cas systems have been widely tested in hippocampal and cortical neurons (Horvath et al., 2017; Ravichandran et al., 2023; Tsunematsu et al., 2017), their application in sensory neurons remains largely unexplored, with only one recent report of successful Cas9 editing in primary human sensory neurons (Palomino et al., 2025). Sensory neurons mediate pain, temperature, and touch sensation and are widely used as a cellular model of nerve regeneration, peripheral neuropathies and spinal cord injury (Maday et al., 2014; Mar et al., 2016; Nascimento et al., 2018). Reliable approaches to manipulate gene expression in these neurons could significantly benefit basic research and therapeutic development, and RNA-targeting CRISPR-Cas systems may be particularly well suited for this purpose. In recent years, several compact RNA-targeting CRISPR-Cas effectors have been characterized and developed, including members of the Cas13 family (type VI) and the Cas7-11 effector (type III-E) (Kato et al., 2022; Özcan et al., 2021; Tong et al., 2022). Although these systems have been successfully used for RNA knockdown in various cell types achieving enhanced performance compared to traditional systems, their performance in sensory neurons is still unknown.

Cas7-11 is a single-protein RNA-targeting CRISPR-Cas effector from the type III-E system (Özcan et al., 2021). It processes its own precursor CRISPR RNA (pre-crRNA) to generate a mature crRNA that directs cleavage of complementary single-stranded RNA at two defined sites via the Cas7.2 and Cas7.3 domains that adopt RRM (RNA recognition motif)-like architecture (Goswami et al., 2022; Kato et al., 2022). Importantly, Cas7-11 mediates precise on-target RNA cleavage without detectable collateral activity and has been shown to achieve efficient on-target knockdown in HEK293 cells and primary human cortical neurons (Kato et al., 2022; Özcan et al., 2021). A truncated Cas7-11 variant, Cas7-11S, was engineered by Kato et al. (2022), substantially reducing protein size while preserving knockdown efficiency.

Cas13 enzymes are single-protein, RNA-guided endonucleases from the type VI system that cleave single-stranded RNA (Abudayyeh et al., 2016; East-Seletsky et al., 2016; Makarova et al., 2025). Cas13 possesses two higher eukaryotes and prokaryotes nucleotide-binding (HEPN) domains, which form an active catalytic site upon target RNA binding and can mediate both target and non-specific (i.e. collateral) RNA degradation (Anantharaman et al., 2013; Konermann et al., 2018). High-fidelity variants of Cas13 have been developed to reduce the collateral RNA cleavage observed with wild type Cas13 (Tong et al., 2022; Wang et al., 2019). The hfCas13d and hfCas13X proteins contain targeted mutations in their HEPN domains that suppress collateral activity while preserving on-target cleavage. These compact variants have shown effective and specific RNA knockdown in mammalian cells (Tong et al., 2022), making them promising tools for gene modulation in sensory neurons. However, neither Cas7-11 nor high-fidelity Cas13 variants have been evaluated for their knockdown efficiency in sensory neurons.

In this study, we evaluated the RNA-targeting CRISPR-Cas systems, Cas7-11S, hfCas13X, and hfCas13d, in primary rat sensory neurons. We assessed their knockdown efficiency and effects on neuronal health through targeting of endogenous mRNA transcripts. Our results reveal clear differences in performance, with Cas7-11S showing toxicity, and hfCas13d emerging as a particularly effective and well-tolerated tool for gene silencing in sensory neurons. These findings provide important insight into the compatibility of RNA-targeting CRISPR-Cas technologies with sensory neurons and offer guidance for the development of effective gene-modulating strategies for neuroscience research and potential therapeutic applications.

## Materials and methods

### Animals

All experiments with animals were performed in compliance with the guidelines for the welfare of experimental animals issued by the Government of New Zealand and were approved by the Animal Ethical Review Committee of the University of Otago AUP22-44.

### Construction of DNA plasmids

The oligonucleotides used in this study are listed in **Supplementary Table 1**. All oligonucleotides used were sourced from Integrated DNA Technologies (USA). All plasmid DNA was purified using Zymo DNA purification kits (Zymo Research). **Supplementary Table 2** lists all the plasmids used in this study. Unless otherwise stated, Gibson assembly, restriction digests, annealing of primers, PCR amplifications, ligations and *Escherichia coli* transformations were performed using standard techniques. DNA from PCR and agarose gels was purified using the Zymoclean Gel DNA recovery kit (Zymo Research). Polymerases, restriction enzymes and T4 ligase were obtained from New England Biolabs or Thermo Fisher.

### Construction of Cas7-11S DNA plasmids

The plasmid pDF0584 huDisCas7-11 S1006-GGGS-D1221 U6-NT guide (LG189), originally described by Kato et al. (2022), was obtained from Addgene (#186993). To enable guide RNA cloning, Gibson assembly and custom-designed primers were used to introduce BbsI restriction sites into the plasmid backbone, generating pDF0584 EF1α-huDisCas7-11 S1006-GGGS-D1221-BbsI (LG208). Six CRISPR-Cas7-11 guides targeting specific sites within the rat MAP2 mRNA (NCBI accession NM_001431749.1), along with two non-targeting control guides, were designed based on previously established protocols and the available online tool for Cas13 by Wessels et al. (2020). MAP2-targeting guides were selected to be nonoverlapping, distributed along the MAP2 transcript, and ranked in the high-scoring quartile (Q4), indicating high predicted knockdown efficacy. The complementary guide sequences were annealed and cloned into LG208 via the introduced BbsI sites. All plasmids were validated by Sanger sequencing conducted by Genetic Analysis Service (GAS) at the University of Otago.

### Construction of hfCas13d DNA plasmids

The Addgene plasmid (#190034) pCBh-NLS_hfCas13d(RfxCas13d_N2V8)_HA_NLS-pA-U6-DR-BpiI-BpiI-pSV40-EGFP-pA-pSV40-mCherry-pA (LG198), originally described by Tong et al. (2022), was used as the backbone for guide cloning. Six CRISPR-hfCas13d guides targeting specific sites within the rat MAP2 mRNA, along with two non-targeting control guides, were designed based on previously established protocols and the available online tool by Wessels et al. (2020). MAP2-targeting guides were selected as described above. The complementary guide sequences were annealed and subcloned into pCBh-NLS_hfCas13d(RfxCas13d_N2V8)_HA_NLS-pA-U6-DR-BpiI-BpiI-pSV40-EGFP-pA-pSV40-mCherry-pA (LG198) via BbsI sites. All resulting constructs were verified by Sanger sequencing conducted by GAS at the University of Otago.

### Construction of hfCas13X DNA plasmids

The Addgene plasmid (#190033) pCBh_NLS_hfCas13X(Cas13X_M17YY)_NLS-pA-U6-DR-BpiI-BpiI-DR-pSV40-EGFP-pA-pSV40-mCherry-pA (LG207), originally described by Tong et al. (2022), was used as the backbone for guide cloning. Six CRISPR-hfCas13X guides targeting specific sites within the rat MAP2 mRNA, along with two non-targeting control guides, were designed based on previously established protocols and the available online tool by Wessels et al. (2020). MAP2-targeting guides were selected as described above. The complementary guide sequences were annealed and subcloned into pCBh_NLS_hfCas13X(Cas13X_M17YY)_NLS-pA-U6-DR-BpiI-BpiI-DR-pSV40-EGFP-pA-pSV40-mCherry-pA (LG207) via BbslI sites. All resulting constructs were verified by Sanger sequencing conducted by GAS at the University of Otago.

### Sensory neuron culture and transfection

Sensory neurons were isolated from adult female Sprague Dawley rats (2-3 months old). The neurons were dissociated with collagenase type IV (Worthington) for 75 minutes and 0.1% trypsin (Sigma-Aldrich) for 15 minutes, followed by gentle trituration with a glass pipette to achieve single-cell suspension. Dissociated neurons were seeded on 18 mm coverslips (Epredia) coated with 20 µg/mL poly-D-lysine (Sigma-Aldrich) and 10 µg/mL laminin (Gibco), and grown in sensory neuron medium containing DMEM (Gibco), 1% fetal bovine serum (In Vitro Technologies), 1% penicillin-streptomycin-fungizone (Gibco). Sensory neurons were kept at 37 °C in 5% CO_2_.

Prior to seeding, dissociated sensory neurons were transfected in suspension using a Neon microporator system (Thermo Fisher). Approximately 1 x 10^5^ cells were transfected per reaction, in a volume of 11 µl with 0.5-1 μg plasmid DNA. Transfected cells were plated and cultured as described above, omitting antibiotics for the first 24h after electroporation. Transfection was performed using the plasmids described in **Supplementary Table 2**.

### Immunofluorescence and confocal imaging

All primary and secondary antibodies used in this study are listed in **Supplementary Table 3**. At five days *in vitro* (DIV) adult rat sensory neurons on glass coverslips were fixed for 15 minutes in 4% paraformaldehyde (Sigma-Aldrich) at room temperature and permeabilized with 0.1% Triton X-100 (Sigma-Aldrich) PBS for 15 minutes. Subsequently they were labelled with primary antibodies in PBS and 10% goat serum (Thermo Fisher) for 2 h followed by secondary antibodies in PBS for 1 h. PBS washes were performed after each antibody incubation. Coverslips were mounted on glass microscope slides (Sail Brand) using Fluorsave mounting medium (Merck Millipore).

Images were captured using an Andor Dragonfly spinning-disk confocal coupled with an inverted Nikon Ti2-E microscope (Nikon), equipped with Nikon CFI Apo TIRF 60× 1.49 NA oil objective (Nikon), Nikon 20× 0.75 NA air objective (Nikon), VC 60× 1.2 NA water objective (Nikon), iXon 888 Andor EMCCD camera, and controlled with Andor Fusion software. All images were taken at the same settings for light and exposure with parameters adjusted to guarantee pixel intensities were below saturation. Montage images were acquired to encapsulate whole neurons, including elaborate axonal elements. All images were scaled processed and analysed using Fiji/ImageJ (NIH) and Adobe IllustratorCS (Adobe Inc.).

### Hoechst and propidium iodide live-stain

5 DIV adult rat sensory neurons cultured on 25 mm glass coverslips (Epredia) were live-stained with propidium iodide (PI) to assess the toxicity of the Cas7-11S expressing plasmids. Rat sensory neurons were incubated with PI (Thermo Fisher, #P1304MP) and Hoechst (Thermo Fisher, #H21486) for 5 minutes at room temperature in the dark. Subsequently, coverslips were placed in a microscope holder (Thermo Fisher) and submerged in imaging media without phenol red (as described by Kühn et al. (2025)) . Live-stained sensory neurons were imaged using an Andor Dragonfly spinning-disk confocal coupled to an inverted Nikon Ti2-E microscope, Nikon VC 60× 1.2 NA water objective (Nikon), iXon 888 Andor EMCCD camera, with environmental control for CO_2_ and temperature. Live imaging of the transfected cells was conducted using Andor Fusion software. All images were taken at the same settings for light and exposure with parameters adjusted so that the pixel intensities were below saturation. All images were processed using Fiji/ImageJ (NIH) and Adobe IllustratorCS (Adobe Inc.).

### Quantification of immunofluorescence in the cell body

MAP2 fluorescence intensity in the cell body of sensory neurons was measured in a standardised manner (area of standard dimension measured at the base of the axon) using Fiji/ImageJ. Background intensity was similarly measured in areas adjacent to the cell body and subtracted from the mean pixel value measured at the cell body giving a final mean fluorescence intensity measurement.

### Quantification of immunofluorescence in the axon

To analyse MAP2 fluorescence intensity in the proximal axon of sensory neurons, the segmented line tool in Fiji/ImageJ was used to draw a line of five pixels wide over the axon from the beginning of the axon towards the end. A line intensity profile from each line was used to analyse the distribution profile of mean fluorescence intensity levels in the proximal axon of MAP2.

### Quantification of propidium iodide

Images were processed using Fiji/ImageJ software and transfected cells were manually assessed for PI fluorescence. Neurons exhibiting PI uptake were classified as PI-positive and the toxicity of the Cas7-11S plasmids was quantified as the percentage of PI-positive neurons relative to the total number of transfected neurons.

### Quantification of cell body circularity

The polygon tool in Fiji/ImageJ was used to trace the cell body of transfected neurons. The measure tool in ImageJ was then used to calculate circularity of the traced cell body as a value between 0 and 1 (value of 1 indicating perfect circularity).

### Statistical analysis

Statistical parameters including the definitions and exact n values (e.g., number of experiments, number of cells), are reported in the figures and figure legends. Data and statistical analysis were performed with Excel (Microsoft) and GraphPad Prism software (GraphPad Software INC). The assumption of data normality was assessed using the D’Agostino-Pearson omnibus test. Statistical analysis includes Ordinary one-way ANOVA with post-hoc Dunnett’s multiple comparisons test and Kruskal-Wallis test with post-hoc Dunn’s multiple comparisons test (not significant is p > 0.05, ^*^p < 0.05, ^**^p < 0.01, ^***^p < 0.001). Graphs were made using GraphPad Prism software. See **Supplementary Table 4** for details on number of experiments, number of cells, type of analysis, statistical test per experiment, and p-values.

## Results

### CRISPR-Cas7-11S is toxic in sensory neurons

To assess the efficiency of RNA-targeting CRISPR effectors in sensory neurons, we first tested the compact CRISPR-Cas7-11S variant to knockdown an endogenous mRNA. Cas7-11S was selected based on its small size, reported on-target activity, lack of nonspecific collateral RNA cleavage, and suitability for *in vivo* applications using viral vectors (Kato et al., 2022). We targeted the rat microtubule associated protein 2 (MAP2), a neuronal cytoskeletal protein involved in cellular architecture and intracellular trafficking (Gumy et al., 2017). In sensory neurons, MAP2 protein localises to the neuronal cell body and is enriched in the initial part of the axon (proximal axon) that extends from the cell body (Gumy et al., 2017). MAP2 transcripts were targeted using six individual guide RNAs (gRNAs) with their targets distributed along the mRNA (**Fig. 1A**). In addition, two non-targeting controls were included (nt1, nt2) to assess potential target-independent effects of Cas7-11S. Each gRNA was expressed under a U6 promoter from a vector also encoding Cas7-11S under an EF1α promoter (Kato et al., 2022). Primary rat adult sensory neurons were co-transfected with individual Cas7-11S/gRNA constructs and a GFP-expressing plasmid to identify transfected neurons. Endogenous MAP2 protein levels in GFP-labelled neurons were quantified by immunofluorescence at five days post-transfection to assess guide-dependent knockdown efficiency (**Fig. 1A**). Control neurons expressing a GFP plasmid alone displayed the expected MAP2 protein distribution, with high fluorescence intensity in the cell body (**Fig. 1B, C**) and a gradual decline in intensity along the proximal axon (**Fig. 1B, D**). In contrast, neurons expressing GFP with the non-targeting gRNAs showed unexpected MAP2 knockdown compared with the GFP control, whereas neurons expressing the targeting gRNAs showed highly variable MAP2 distribution and fluorescence intensity within the cell body and proximal axon (**Fig. 1B, C, D, E**).

**Figure 1.**
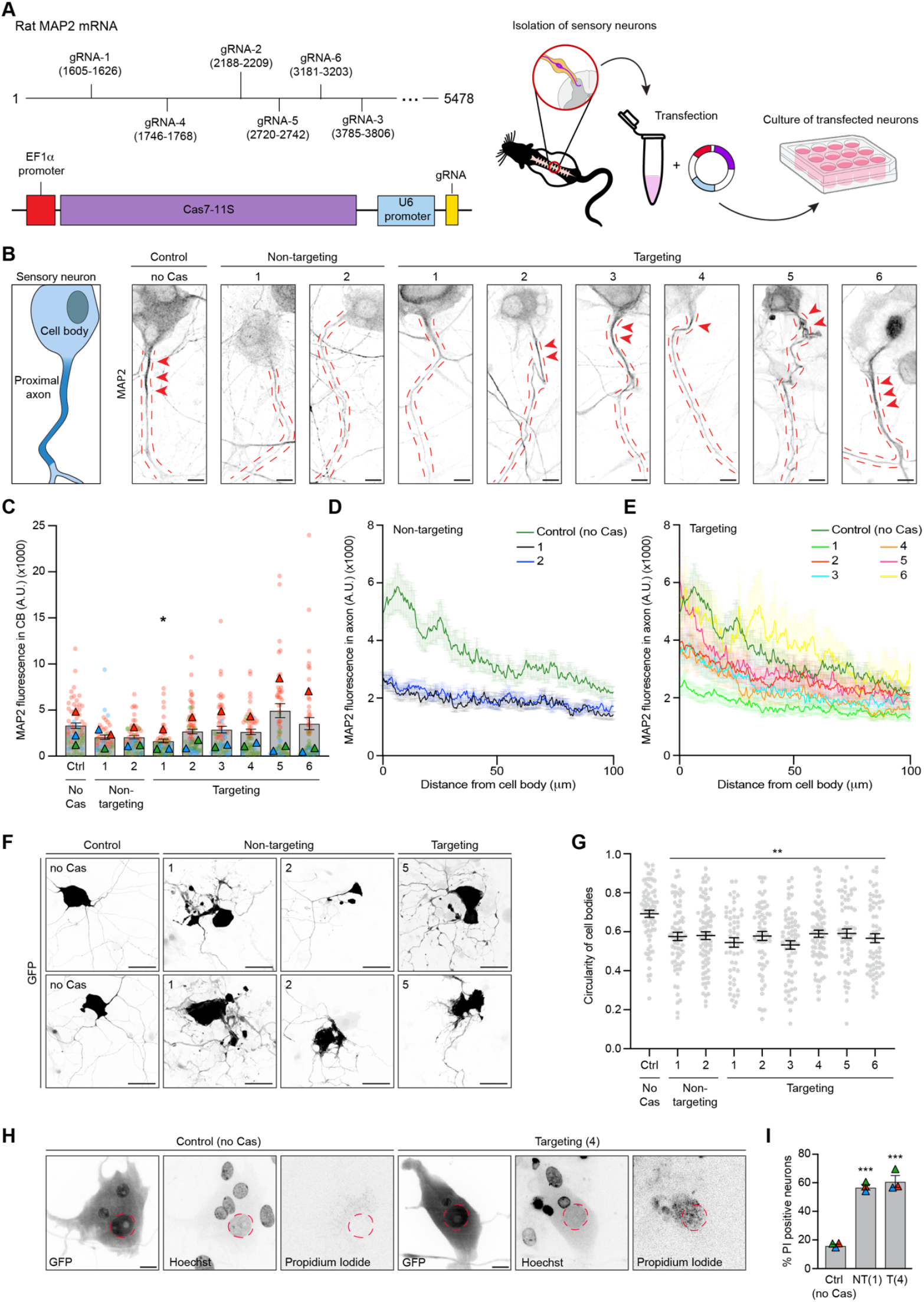
CRISPR-Cas7-11S is toxic in sensory neurons. (**A**) Schematics showing CRISPR-Cas7-11S guide RNA (gRNA) targeting positions within the MAP2 mRNA, the CRISPR-Cas7-11S plasmid and the experimental set-up. (**B**) Representative images of adult sensory neurons (5 DIV) immunostained for endogenous MAP2 protein under the indicated conditions. Dashed red lines outline the axon. Arrowheads indicate MAP2 enrichment in the proximal axon. Scale bar 10 μm. (**C**) Quantification of MAP2 fluorescence intensity in the cell body of sensory neurons (5 DIV) under the indicated conditions. *N*= 3 independent experiments with at least 45 neurons per condition (A.U. arbitrary units). Kruskal-Wallis test with post-hoc Dunn’s multiple comparisons test. (**D, E**) Average MAP2 fluorescence intensity along the proximal axon of sensory neurons (5 DIV) under the indicated conditions *N*= 3 independent experiments with at least 34 neurons per condition. (**F**) Representative images showing sensory neuron (5 DIV) cell body morphology under the indicated conditions. Scale bar 50 μm. (**G**) Quantification of sensory neuron cell body circularity under the indicated conditions. *N*= 3 independent experiments with at least 53 neurons per condition. Kruskal-Wallis test with post-hoc Dunn’s multiple comparisons test. (**H**) Representative images of sensory neurons (5 DIV) under the indicated conditions live-stained with Hoechst nucleic acid dye and propidium iodide. Red dashed circles indicate the nucleus of the transfected neuron. Scale bar 10 μm. (**I**) Quantification of the percentage of propidium iodide (PI)-positive sensory neurons under the indicated conditions. *N*= 3 independent experiments with at least 44 cells per condition. Ordinary one-way ANOVA with post-hoc Dunnett’s multiple comparisons test. All data are presented as mean values ± SEM. ^*^p<0.05, ^**^p<0.01, ^***^p<0.001.

We hypothesized that the variable knockdown of MAP2 was due to cellular stress caused by Cas7-11S. To test this, we examined cell viability and morphology of sensory neurons transfected with Cas7-11S constructs. Control GFP-transfected sensory neurons displayed normal morphology with rounded cell bodies and elaborate axonal architecture (**Fig. 1F**) (Scott, 1977). In contrast, sensory neurons expressing any of the Cas7-11S constructs tested, including control or targeting gRNAs, displayed abnormal morphology, with malformed cell bodies, swellings and bulbs at the axon tips (**Fig. 1F**). Since alterations in cell body shape indicate cellular stress, we next analysed cell body circularity. Notably, all Cas7-11S constructs irrespective of gRNA significantly decreased cell body circularity compared with the GFP control, suggesting a morphological shift toward a less rounded phenotype (**Fig. 1G**), consistent with sensory neurons undergoing cellular damage (Charras, 2008). To more directly assess Cas7-11S toxicity, we performed propidium iodide (PI) live-staining. The fraction of PI-positive nuclei, indicative of membrane-compromised (i.e. dying or dead) neurons, was significantly increased in all Cas7-11S conditions examined compared with the GFP control (**Fig. 1H, I**). In summary, Cas7-11S expression in sensory neurons induces cytotoxic effects that elicit altered neuronal morphology and increased cell death.

### CRISPR-hfCas13X fails to knockdown MAP2 in sensory neurons

Given the toxicity observed with Cas7-11S, we next assessed whether an alternative RNA-targeting system could provide more efficient gene silencing in sensory neurons. The hfCas13X protein, a compact high-fidelity Cas13 variant engineered by Tong et al. (2022), showed on-target degradation of RNAs and minimal collateral effects in HEK293 cells and *in vivo* in mice. We therefore tested hfCas13X for its guide-dependent knockdown efficiency in sensory neurons. The six MAP2-targeting gRNAs and two non-targeting controls (nt1, nt2) from our previous experiments were inserted into a vector expressing GFP under the SV40 promoter and hfCas13X under a CBh promoter and individual gRNAs under a U6 promoter (Tong et al., 2022) (**Fig. 2A**). Primary rat sensory neurons were transfected with individual hfCas13X/gRNA constructs, and MAP2 protein levels were assessed by immunofluorescence five days post-transfection. Among the six MAP2-targeting guides, only gRNA-2 and gRNA-4 produced a significant reduction in MAP2 cell body fluorescence intensity compared with nt2 (**Fig. 2B, C**), while the other MAP2-targeting guides showed MAP2 levels in the cell body comparable to controls, indicating that hfCas13X did not reliably knock down MAP2 in these neurons. Analysis of MAP2 fluorescence in the proximal axon similarly showed a reduction in axonal MAP2 fluorescence intensity for gRNA-2 and gRNA-4, while the other MAP2-targeting guides showed axonal MAP2 levels comparable to controls (**Fig. 2D, E**). Expression of hfCas13X did not induce toxicity, since neurons retained normal morphology, and cell body circularity was unchanged compared to controls (**Fig. 2F, G**). Overall, these findings show that while hfCas13X expression from these plasmids is well tolerated in sensory neurons, it fails to achieve robust and reproducible knockdown of endogenous MAP2.

**Figure 2.**
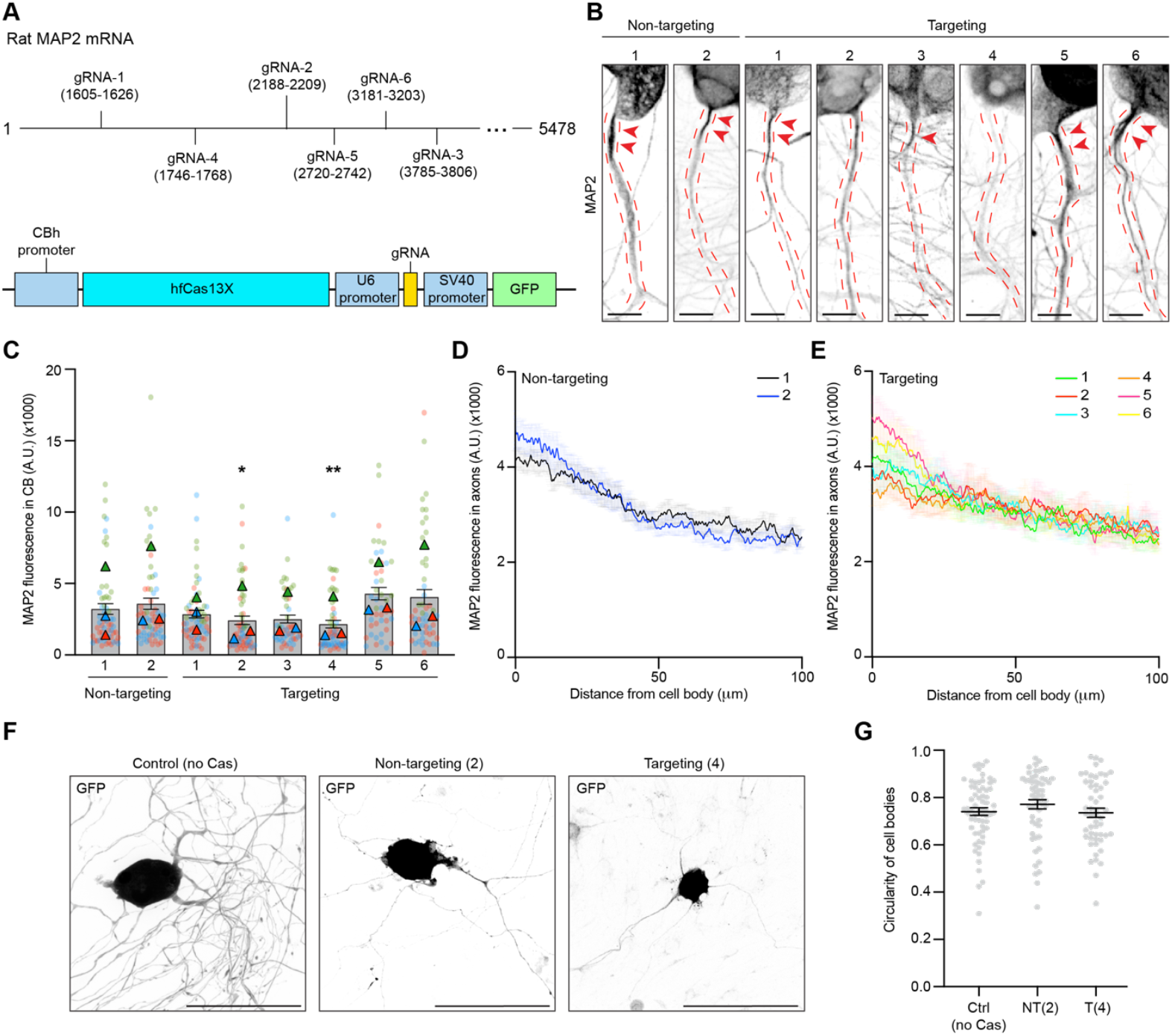
CRISPR-hfCas13X fails to knockdown MAP2 in sensory neurons. (**A**) Schematics showing CRISPR-hfCas13X guide RNA (gRNA) targeting positions within the MAP2 mRNA and the CRISPR-hfCas13X plasmid. (**B**) Representative images of adult sensory neurons (5 DIV) immunostained for endogenous MAP2 protein under the indicated conditions. Dashed red lines outline the axon. Arrowheads indicate MAP2 enrichment in the proximal axon. Scale bar 10 μm. (**C**) Quantification of MAP2 fluorescence intensity in the cell body of sensory neurons (5 DIV) under the indicated conditions. *N*= 3 independent experiments with at least 47 neurons per condition (A.U. arbitrary units). Kruskal-Wallis test with post-hoc Dunn’s multiple comparisons test. (**D, E**) Average MAP2 fluorescence intensity along the proximal axon of sensory neurons (5 DIV) under the indicated conditions. *N*= 3 independent experiments with at least 54 neurons per condition. (**F**) Representative images showing sensory neuron (5 DIV) cell body morphology under the indicated conditions. Scale bar 100 μm. (**G**) Quantification of sensory neuron cell body circularity under the indicated conditions. *N*= 3 independent experiments with at least 52 neurons per condition. Kruskal-Wallis test with post-hoc Dunn’s multiple comparisons test. All data are presented as mean values ± SEM. ^*^p<0.05, ^**^p<0.01.

### CRISPR-hfCas13d efficiently silences MAP2 in sensory neurons

Given the limited knockdown efficiency achieved with hfCas13X, we next evaluated an alternative Cas13 protein, hfCas13d, to determine whether it provides more effective gene silencing in sensory neurons. The hfCas13d protein is another high-fidelity RNA-targeting variant engineered by Tong et al. (2022) and was tested here using the same six gRNAs targeting along the rat MAP2 transcript (**Fig. 3A**). In addition, two non-targeting controls were included (nt1, nt2). All guides were cloned under the U6 promoter into a vector encoding GFP under the SV40 promoter and hfCas13d driven by the CBh promoter (Tong et al., 2022) (**Fig. 3A**). Primary rat sensory neurons were transfected with individual hfCas13d/gRNA constructs, and MAP2 protein levels were assessed by immunofluorescence five days post-transfection. Representative images show a marked reduction of MAP2 fluorescence in the cell body and the proximal axon for all gRNAs compared to the controls (**Fig. 3B**). Quantitative analysis of MAP2 fluorescence intensity in the cell body confirmed significant reductions in MAP2 fluorescence intensity for all gRNAs compared with nt1/2, with on average 50-60% reductions of MAP2 irrespective of the guide used (**Fig. 3C**). Analysis of MAP2 fluorescence in the proximal axon likewise showed reduced intensity, indicating that hfCas13d-mediated MAP2 knockdown decreases MAP2 levels throughout the cell body and proximal axon (**Fig. 3D, E**). Importantly, hfCas13d expression did not induce toxicity, since neuronal morphology and cell body circularity remained unchanged compared with the controls (**Fig. 3F, G**). These findings demonstrate that CRISPR-hfCas13d can be programmed with specific gRNAs to effectively silence endogenous MAP2 in sensory neurons, with knockdown observed in both the cell body and proximal axon. These results establish CRISPR-hfCas13d as a reliable tool for targeted gene silencing in sensory neurons.

**Figure 3.**
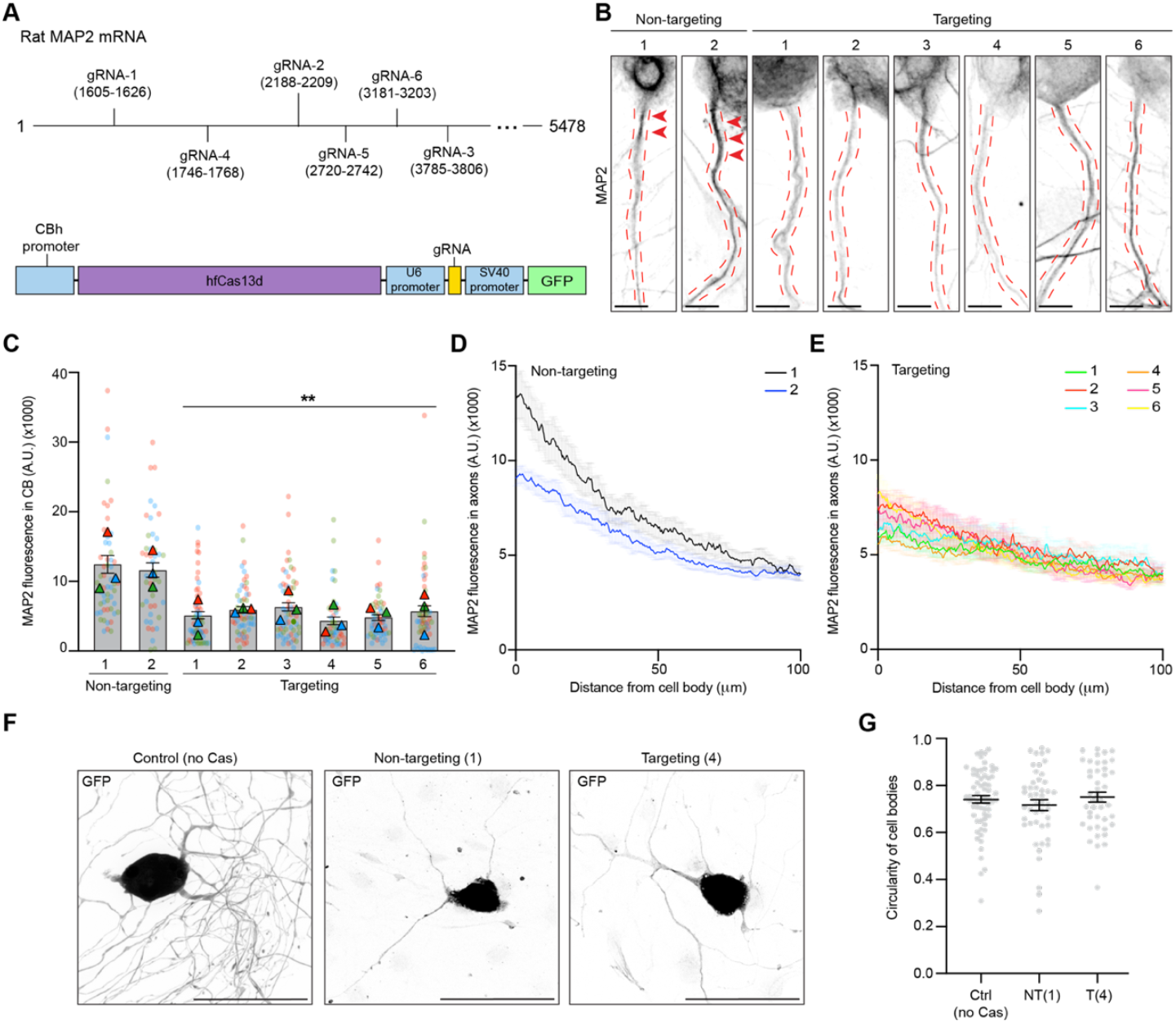
CRISPR-hfCas13d efficiently silences MAP2 in sensory neurons. (**A**) Schematics showing CRISPR-hfCas13d guide RNA (gRNA) targeting positions within the MAP2 mRNA and the CRISPR-hfCas13d plasmid. (**B**) Representative images of adult sensory neurons (5 DIV) immunostained for endogenous MAP2 protein under the indicated conditions. Dashed red lines outline the axon. Arrowheads indicate MAP2 enrichment in the proximal axon. Scale bar 10 μm. (**C**) Quantification of MAP2 fluorescence intensity in the cell body of sensory neurons (5 DIV) under the indicated conditions. *N*= 3 independent experiments with at least 46 neurons per condition (A.U. arbitrary units). Kruskal-Wallis test with post-hoc Dunn’s multiple comparisons test. (**D, E**) Average MAP2 fluorescence intensity along the proximal axon of sensory neurons (5 DIV) under the indicated conditions. *N*= 3 independent experiments with at least 73 neurons per condition. (**F**) Representative images showing sensory neuron (5 DIV) cell body morphology under the indicated conditions. Scale bar 100 μm. (**G**) Quantification of sensory neuron cell body circularity under the indicated conditions. *N*= 3 independent experiments with at least 43 neurons per condition. Kruskal-Wallis test with post-hoc Dunn’s multiple comparisons test. All data are presented as mean values ± SEM. ^**^p<0.01.

## Discussion

In this study, we compared the RNA-targeting CRISPR effectors Cas7-11S, hfCas13X, and hfCas13d, for their ability to knock down endogenous MAP2 in sensory neurons. Sensory neurons are an established cellular model for studying axon growth and regeneration, pain mechanisms, and neuron–environment interactions (Maday et al., 2014; Mar et al., 2016; Nascimento et al., 2018). Reliable gene expression tools for this cell type could therefore benefit both basic and translational research. Our findings highlight hfCas13d as the most reliable RNA-targeting CRISPR tool for sensory neurons, achieving efficient and well-tolerated knockdown, while hfCas13X showed limited knockdown and Cas7-11S caused inconsistent knockdown with cytotoxic effects.

Despite promising reports of on-target cleavage with minimal collateral effects (Kato et al., 2022; Özcan et al., 2021), expression of Cas7-11S in sensory neurons produced highly variable MAP2 levels and pronounced toxicity, regardless of whether MAP2-targeting or non-targeting gRNAs were used. Our findings are consistent with reports describing low knockdown efficiency and non-specific activity of Cas7-11 in HEK293 and Neuro2A cells (McCallister et al., 2023; Zeballos et al., 2023). The observed toxicity in our cellular model may reflect sensory neuron-specific sensitivity to Cas7-11S expression or mechanisms.

Although hfCas13d and hfCas13X have been shown to mediate effective and specific RNA knockdown in mammalian cells (Tong et al., 2022), hfCas13X expression in sensory neurons produced only modest MAP2 knockdown despite no detectable toxicity. This limited activity aligns with previous reports indicating that hfCas13X, while reducing collateral activity compared with wild type Cas13X, also exhibited reduced on-target cleavage efficiency (Liu et al., 2024; Tong et al., 2022). Recent work demonstrating improved on-target activity through strategic modifications of its associated crRNA (Liu et al., 2024) suggests that optimized use of hfCas13X could still represent a suitable approach for gene silencing in sensory neurons and should be tested in future studies.

hfCas13d achieved robust MAP2 knockdown without affecting neuronal morphology. This finding is consistent with previous studies reporting efficient hfCas13d-mediated endogenous mRNA knockdown across various cellular models without detectable toxicity (Chen et al., 2025; McCallister et al., 2023; Zeballos et al., 2023) and further supports evidence that hfCas13d outperforms hfCas13X (Chen et al., 2025). Although Cas13 systems can induce collateral RNA cleavage upon target RNA binding (Abudayyeh et al., 2016; East-Seletsky et al., 2016), the use of high-fidelity Cas13d likely mitigates this effect (Back et al., 2025) as we observed no cytotoxicity in our experiments.

Overall, our findings indicate that hfCas13d is currently the most reliable RNA-targeting CRISPR tool for gene silencing in sensory neurons, whereas Cas7-11S and hfCas13X may require further engineering or optimization to achieve comparable performance in this context. Notably, hfCas13d-mediated knockdown in other types of neuronal cells is already being applied in several *in vivo* studies, including models of Amyotrophic Lateral Sclerosis (ALS) (McCallister et al., 2023) and ocular hypertension (Chen et al., 2025). Our demonstration of its compatibility with sensory neurons further reinforces hfCas13d as a promising RNA-targeting CRISPR-Cas enzyme for future therapeutic applications.

## Acknowledgements

Microscopy was performed in the Confocal Microscopy Unit, Research Infrastructure Centre at the University of Otago, Dunedin, New Zealand

pDF0584 huDisCas7-11 S1006-GGGS-D1221 U6-NT guide was a gift from Omar Abudayyeh & Jonathan Gootenberg (Addgene plasmid # 186993; http://n2t.net/addgene:186993; RRID:Addgene_186993). pCBh-NLS_hfCas13d(RfxCas13d_N2V8)_HA_NLS-pA-U6-DR-BpiI-BpiI-pSV40-EGFP-pA-pSV40-mCherry-pA was a gift from Huawei Tong (Addgene plasmid # 190034 ; http://n2t.net/addgene:190034; RRID:Addgene_190034). pCBh_NLS_hfCas13X(Cas13X_M17YY)_NLS-pA-U6-DR-BpiI-BpiI-DR-pSV40-EGFP-pA-pSV40-mCherry-pA was a gift from Huawei Tong (Addgene plasmid # 190033; http://n2t.net/addgene:190033; RRID:Addgene_190033)

## Author contributions

**AM:** methodology, validation, formal analysis, investigation, writing – original draft, visualisation. **MP:** methodology, validation, formal analysis, investigation, visualisation. **NB**: methodology, investigation, supervision. **DM-M:** methodology, investigation, supervision. **GS:** investigation. **PF:** conceptualisation, resources, writing - original draft, supervision, funding acquisition. **RF:** conceptualisation, investigation, writing - original draft, supervision. **LFG:** conceptualisation, resources, writing - original draft, supervision, project administration, funding acquisition.

## Statements and Declarations

### Consent to participate

Not applicable

### Consent for publication

Not applicable

### Funding statement

The authors disclose receipt of the following financial support for the research, authorship, and/or publication of this article: This work was supported by the Health Research Council of New Zealand [grant number 21-080]. PCF, NB, DM-M and RDF were supported by Bioprotection Aotearoa (Tertiary Education Commission, NZ) and PCF was supported by a James Cook Research Fellowship (Royal Society of New Zealand, Te Apārangi). AM and DM-M were supported by University of Otago Doctoral scholarships. NB was supported by a Health Sciences Career Development Postdoctoral Fellowship (University of Otago). GS was supported by a School of Biomedical Sciences Summer Research Scholarship.

---

**Supplementary Table 1:**
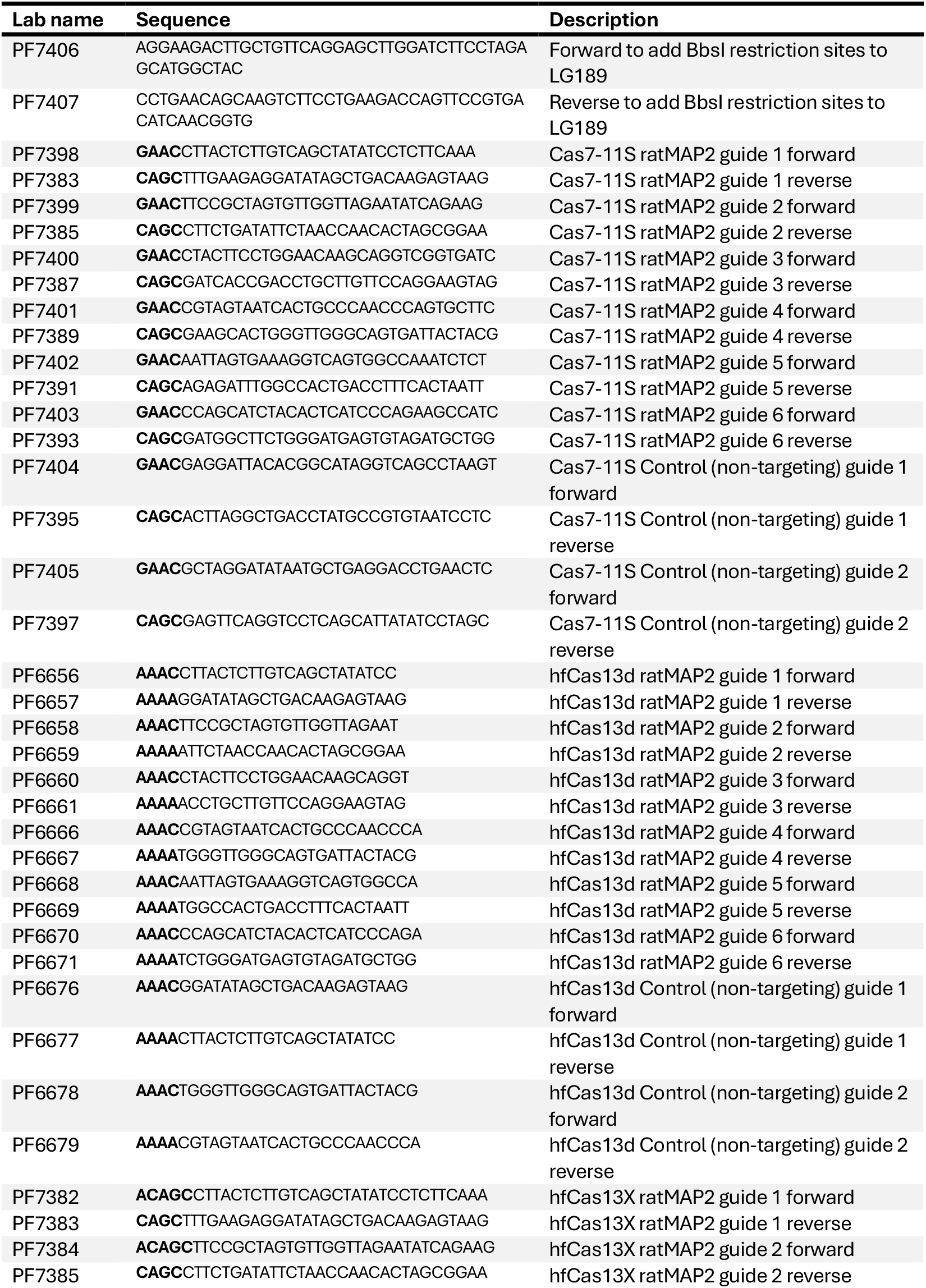
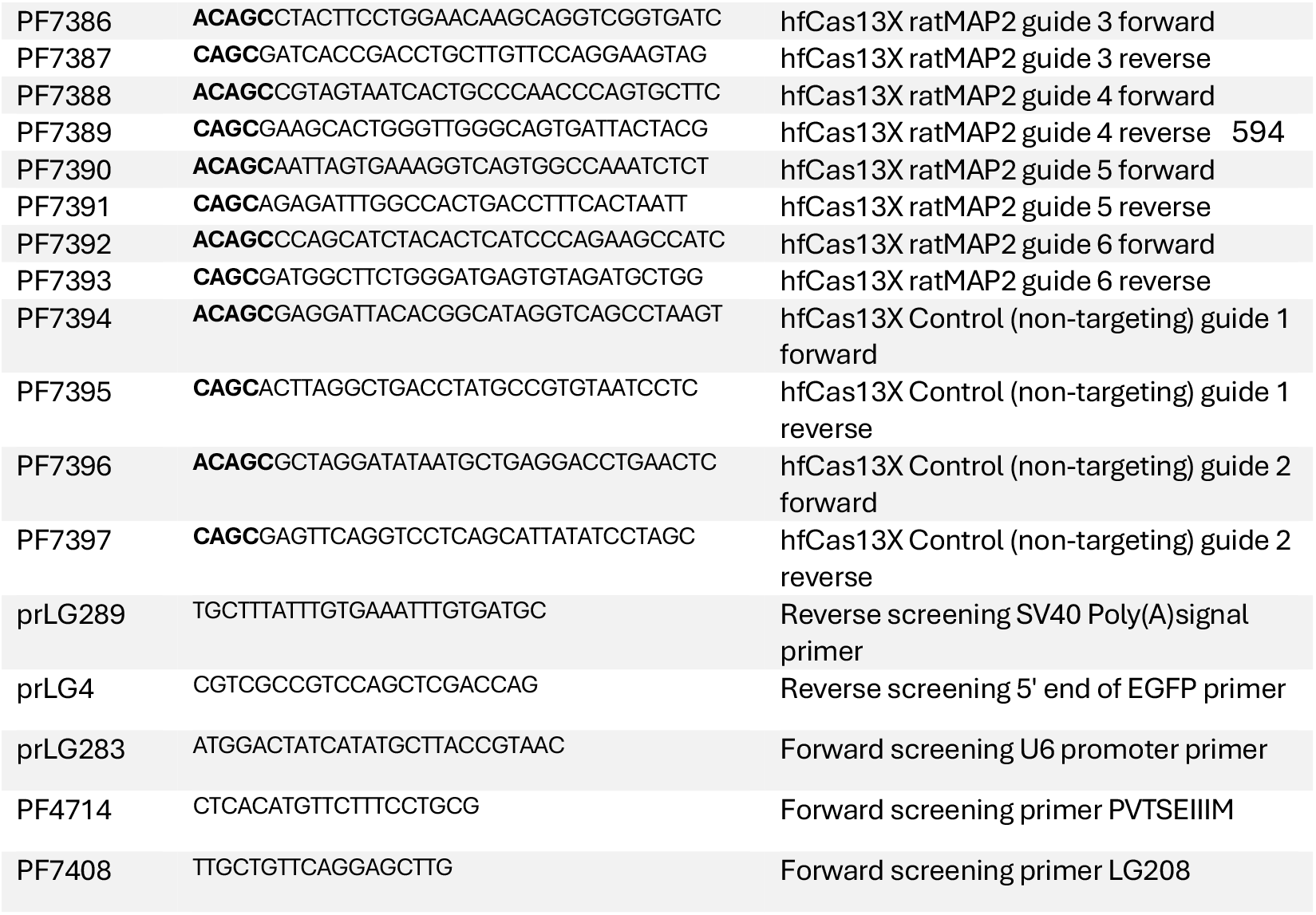
Oligonucleotides used in this study.

**Supplementary Table 2:**
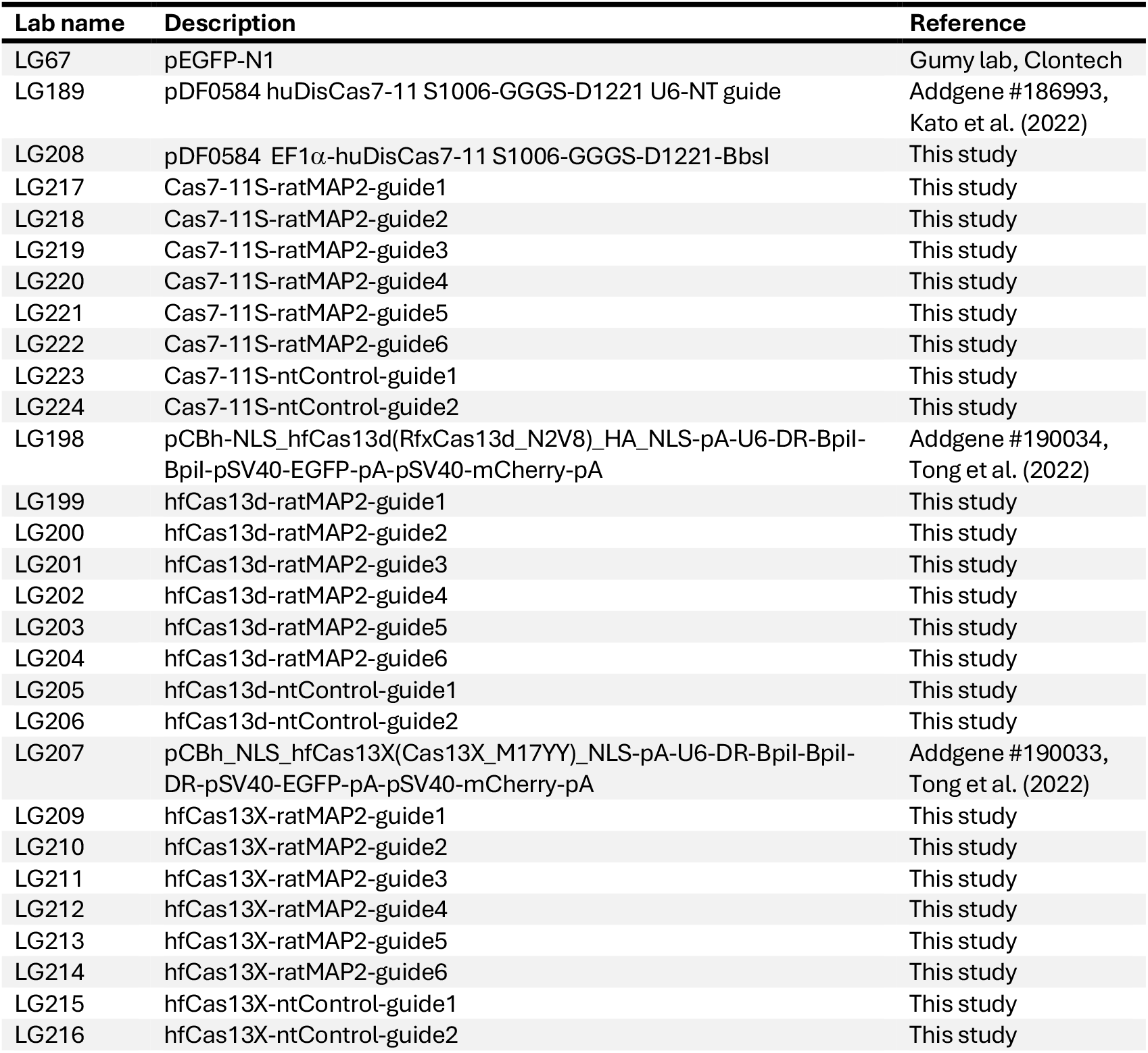
Plasmids used in this study.

**Supplementary Table 3:**
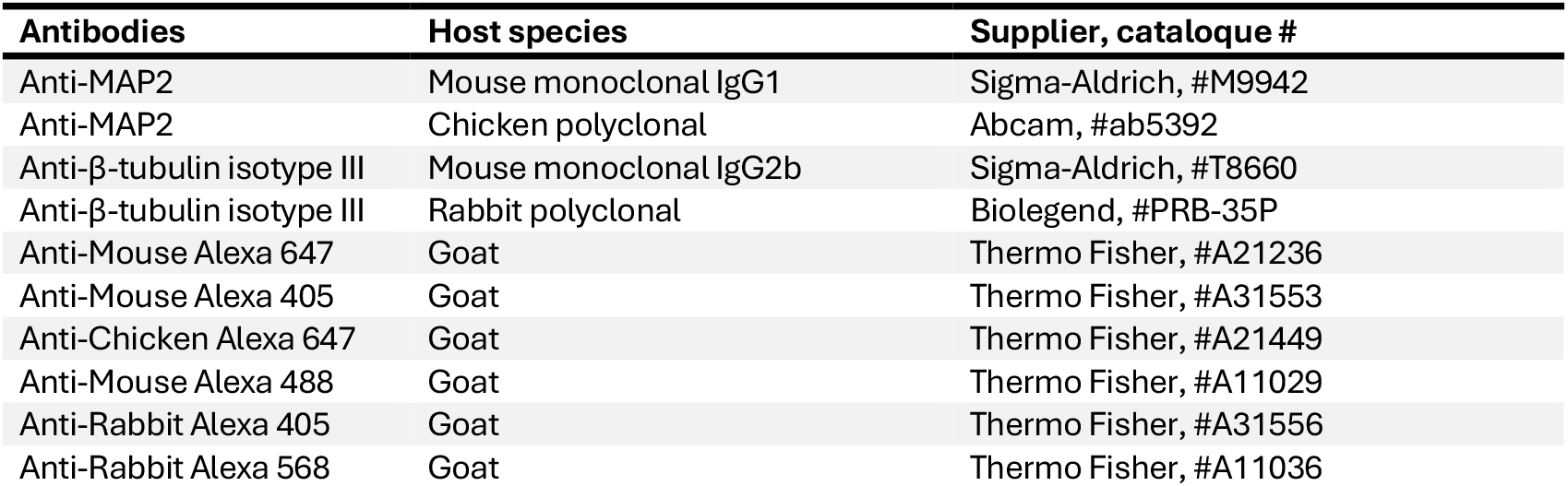
Antibodies used in this study.

**Supplementary Table 4:**
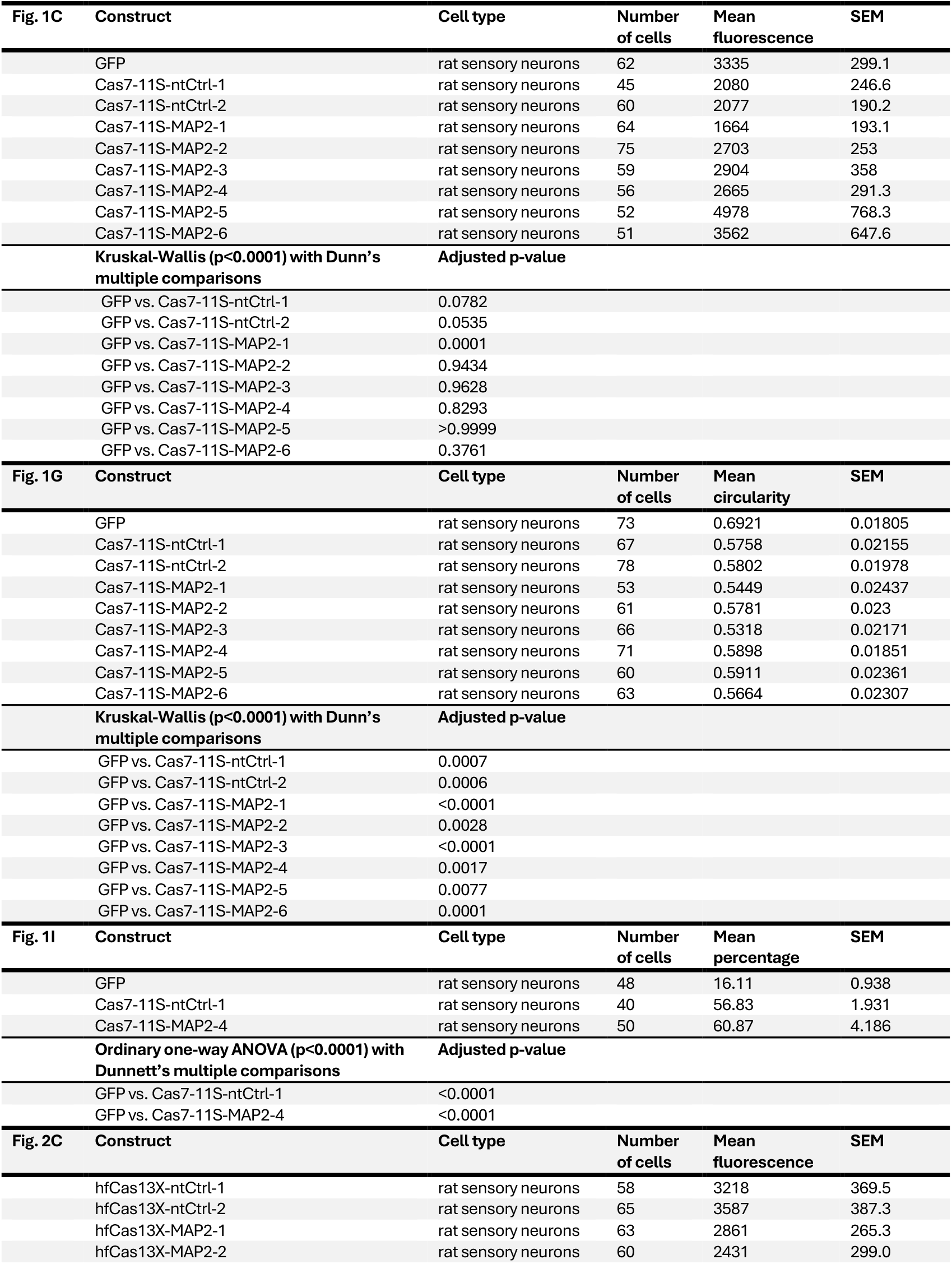
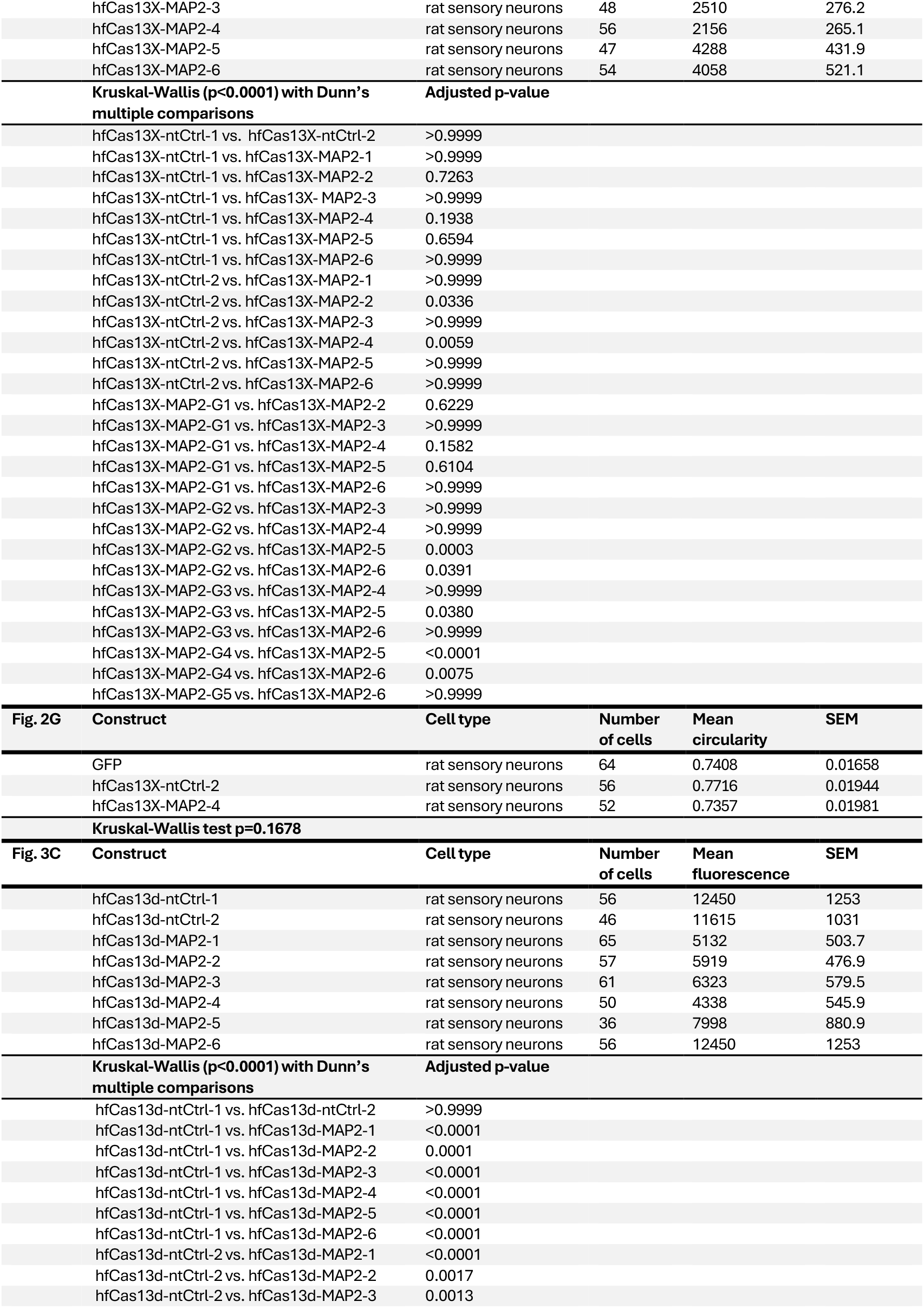
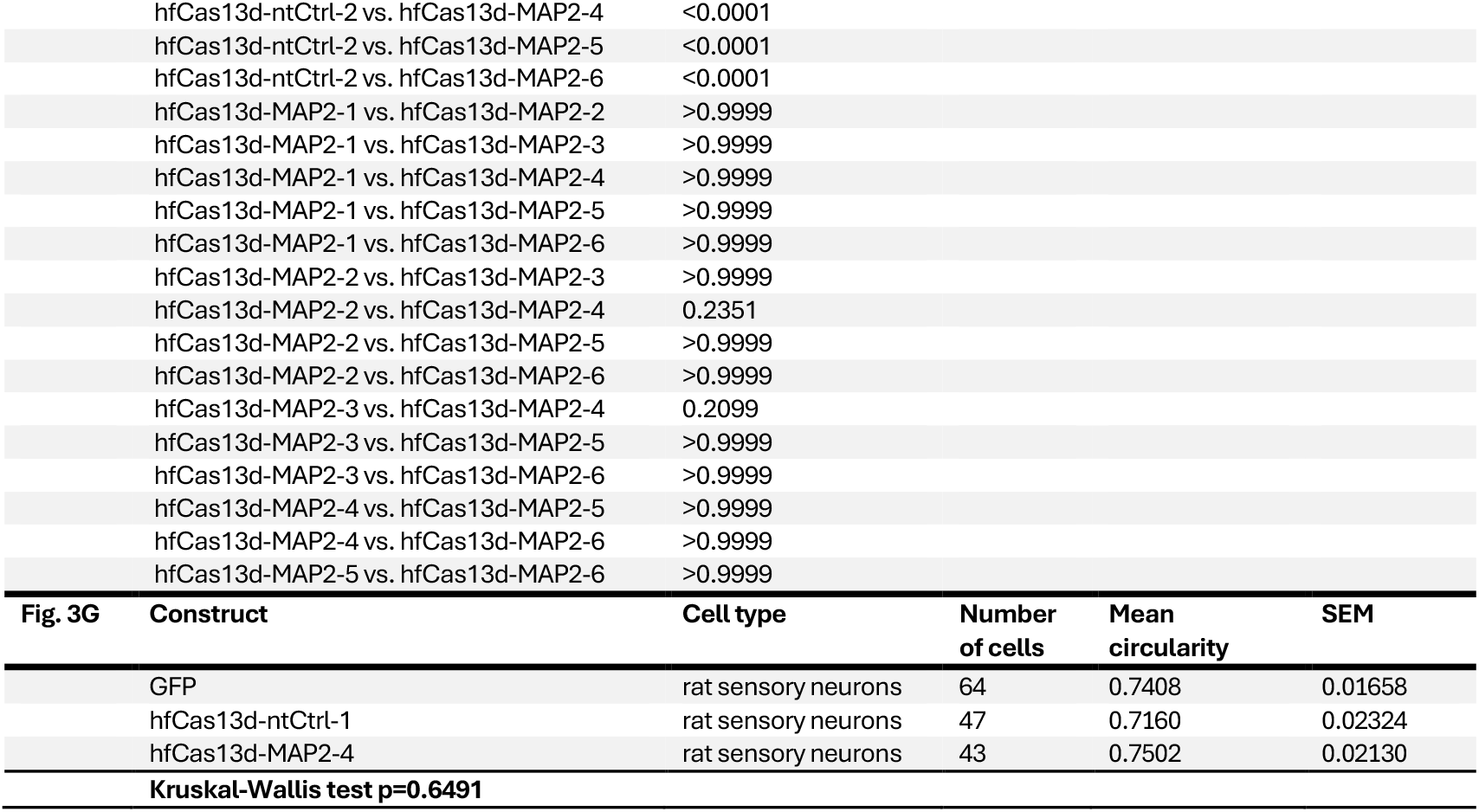
Number of cells, distributions, deviations, statistical tests and p-values for all experiments performed in this study.

## Notes

### Competing Interest Statement

The authors have declared no competing interest.

